# Mechanism of Ribosome Stalling by the AMD1 C-terminal Tail Arrest Peptide

**DOI:** 10.1101/2025.11.16.688537

**Authors:** Emir Maldosevic, Fabio S. Boiocchi, Michal I. Swirski, Kyle A. Meiklejohn, Martina M. Yordanova, Pavel V. Baranov, Ahmad Jomaa

## Abstract

*AMD1* encodes Adenosylmethionine decarboxylase 1 (AMD1), a key enzyme required for polyamine biosynthesis. A subset of ribosomes translating the *AMD1* coding sequence (CDS) read through the stop codon and pause at the next in-frame stop codon 384 nucleotides downstream. The resulting C-terminal extension (C-tail) is universally conserved across all vertebrates, implying that its molecular function is critical to their fitness. Despite growing evidence that such cis-acting elements regulate translation of their genes, the molecular mechanism by which the C-tail mediates ribosome stalling remains unclear. Here, we determined the structure of the ribosome nascent chain complex paused by the AMD1 C-tail which traps eukaryotic release factor 1 (eRF1) with the ATP-binding cassette sub-family E member 1 (ABCE1). The nascent chain forms a molecular clamp that positions an arginine finger in the peptidyl-transferase center, occluding the accommodation of the eRF1 GGQ motif thereby hampering translation termination. Analysis of aggregated ribosome profiling data revealed several genes with a pattern of stop codon readthrough followed by ribosome stalling at a specific location, suggesting that regulatory readthrough-stall mechanisms may not be limited to *AMD1*.

## Introduction

Cells respond to environmental cues and shifting metabolic demands by regulating gene expression to adapt and ensure survival. Regulation can occur at the level of protein synthesis during different steps of translation, including elongation and termination (*1*–*10*). Emerging evidence indicates that ribosomes actively control protein synthesis by sensing elements encoded in cellular mRNAs and nascent chains as they pass through the mRNA channel or polypeptide exit tunnel, respectively. Programmed ribosome pausing along these sequence elements modulates gene expression (*11*–*14*), regulates metabolism (*15*–*18*) and enables cells to overcome intracellular stress (*19*–*22*).

Polyamines are ubiquitous polycationic molecules that bind to negatively charged nucleic acids, regulate their metabolism, and are essential for cell growth and proliferation (*23*–*25*). At the level of protein synthesis, polyamines regulate translation efficiency and fidelity (*26, 27*). Intriguingly, nascent peptides encoded by key polyamine metabolism enzyme transcripts have emerged with a role in sensing polyamine levels from within the ribosomal tunnel, enabling mechanisms such as ribosome stalling (*16, 28, 29*) and frameshifting (*30, 31*).

Adenosylmethionine decarboxylase 1 (AMD1, also known as AdoMetDC), is one such enzyme essential for both spermine and spermidine biosynthesis (*32*). The conserved stalling peptide, MAGDIS, encoded by a regulatory translon (*33*) located close to the 5’ end of the *AMD1* mRNA controls AMD1 enzyme levels depending on intracellular spermidine concentrations (*28, 34, 35*). Under elevated spermidine conditions, ribosomes pause at the stop codon of the MAGDIS-encoding translon. The paused ribosome would then sterically block assembly of preinitiation complexes at *AMD1* mRNA preventing translation initiation at the downstream start codon in the *AMD1* coding sequence (CDS), thereby reducing AMD1 synthesis.

A recent study used ribosome profiling to identify a second highly conserved stalling site downstream of *AMD1* CDS (*36*). During translation of the coding sequence a small subset of ribosomes read through the stop codon (*36, 37*) and lead to the incorporation of a 127 amino acid long C-tail into the enzyme. Ribosomes will then pause at a downstream in-frame stop codon exposing the AMD1 C-tail while it remains tethered to the ribosome. Addition of this C-tail leads to a reduction in the levels of the extended protein (*36, 38*). Despite growing evidence that cis-acting elements regulate translation of their gene (*1, 2, 14*), the precise regulatory elements employed by the AMD1 C-tail remain unclear. Furthermore, it is not understood why nascent chains fail to release from ribosomes stalled at a stop codon of the AMD1 C-tail extension, suggesting that translation termination is impaired.

## Results

### eRF1 and ABCE1 are trapped on ribosomes translating the AMD1 C-tail

Analysis of publicly available ribosome profiling data in *AMD1* locus using RiboSeq.Org tools (*39*) showed peaks of high ribosome density both upstream and downstream of the *AMD1* CDS (*36*). A prominent peak, 384 nucleotides downstream of the coding sequence, corresponds to ribosomes stalled at a stop codon following the translation of the AMD1 C-tail (Fig. 1A-B, ‘Stalled Ribosomes’). As the number of vertebrates with available ribosome profiling data have increased since the original publication (*36*), we have expanded the analysis to these species. Peaks of ribosomal footprints corresponding to the stalling location in human *AMD1* mRNA are evident in most species with available data despite varying coverage and quality of the data (fig. S1). This suggests that not only C-tail extension, but also ribosome stalling mediated by it, are universally conserved across vertebrates.

**Fig. 1.**
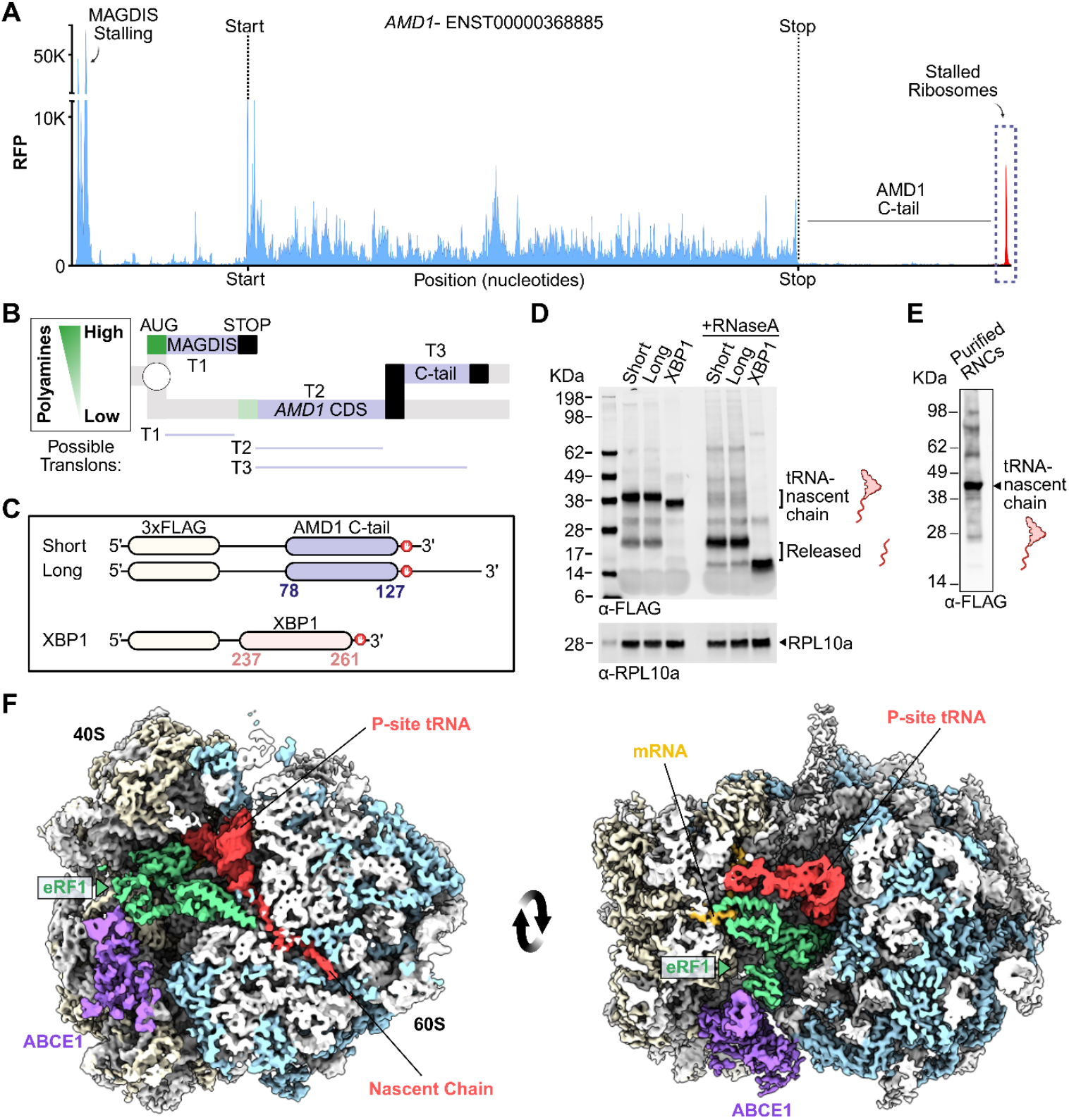
Cryo-EM structure of ribosomes stalled on the 3’UTR of the AMD1 mRNA. A) Ribosome profile of the *AMD1* transcript highlighting the observed ribosome pausing events in the 5’ and 3’ UTRs generated with ribocrypt.org. B) Ribosome decision graph showing translons (T1, T2 and T3) organization of the *AMD1* mRNA illustrating polyamine-dependent control of *AMD1* CDS translation. C) Schematic diagram for the mRNA used to stall ribosomes with the AMD1 C-tail (AAs 78-127) and with XBP1 (AAs 237-261) as a control. D) In vitro translation reactions of the indicated mRNAs from (C). E) Western blot of purified ribosomes stalled at the C-tail of AMD1. F) Cryo-EM density map of the ternary complex filtered to 4Å for visualization. Cross sections of the map shows that the P-site tRNA and nascent chain remain attached to ribosomes with ABCE1 and eRF1 bound. Colors: grey-rRNA; blue- large subunit r-proteins; beige- small subunit r-proteins; yellow- mRNA; purple- ABCE1; green- eRF1; red- tRNA-NC.

To reconstitute this pausing event, we generated constructs encoding the C-tail of AMD1 fused to an N-terminal 3X-FLAG via a flexible linker and monitored ribosome stalling using an *in vitro* translation system in rabbit reticulocyte lysates (RRL) (Fig. 1C-D). Immunoblot analysis demonstrated that ribosomes were stalled with a nascent chain still bound to a P-site tRNA, as indicated by a single band at ∼45 kDa, the combined molecular weights of the nascent chain (20 kDa) and tRNA (25 kDa). Treating the reactions with RNase A released nascent chains from the tRNA indicated by a shift in the nascent chain band size to ∼20 kDa (Fig. 1D). Further extending the mRNA construct by an additional 135 bases downstream of the C-tail stop codon did not shift the molecular weight of the stalled product when compared to the shorter construct (Fig. 1D). These results indicate that pausing occurs at the stop codon of *AMD1* extended translon without the release of the nascent chain consistent with previous results (*36*).

Next, we purified the AMD1 C-tail ribosome nascent chain complex (RNC_AMD1C_) and determined its structure using cryogenic electron microscopy (cryo-EM) (Fig. 1E-F). An initial 2D classification followed by a subsequent 3D classification of the particles were conducted to remove non-ribosomal or heterogeneous particles in different translation states (fig. S2A). This approach yielded a cryo-EM map of the stalled 80S ribosome that displayed strong density for the P-site tRNA and a nascent chain present in the polypeptide exit tunnel (fig. S2A, red). The density of the nascent chain can be unambiguously assigned to the AMD1 arresting peptide tethered to the P-site tRNA within the ribosome. Furthermore, two additional bound factors were observed at the interface of the ribosomal subunits (40S and 60S) (fig. S2B-C, purple). One of the bound factors is the release factor eRF1 resolved interacting with mRNA in the A-site (Fig. 1F, fig. S3A-D, green and yellow). The second density corresponded to the recycling factor ABCE1 (Fig. 1F, fig. S3E, purple) and together these components form the ternary ABCE1:eRF1:RNC_AMD1C_complex (Supplementary Movie 1, Table S1).

### The GGQ motif of eRF1 is sequestered in an inactive conformation

Translation termination is a multi-step process that requires stop codon recognition and hydrolysis of the peptidyl-tRNA ester bond to release nascent proteins from the ribosome (*40, 41*). The N-domain of eRF1 recognizes stop codons in the A-site of the ribosome (Fig. 2A, ‘N’), inducing a conformational change within eRF1 that extends its M-domain (Fig 2A, ‘M’) toward the peptidyl transferase center (PTC) to promote peptide hydrolysis (*6, 42*–*45*). The M-domain contains a universally conserved GGQ motif located in a flexible loop adjacent to α-helix 5 (Fig. 2A, ‘αh5’) which is used to facilitate the hydrolysis reaction *via* a methylated Gln 185 residue (*46*–*48*). Previous studies have shown that ABCE1 co-purifies with eRF1 on ribosomes, indicating that eRF1 serves as a platform for ABCE1 binding (*43, 44*). This enables ABCE1 to split and recycle ribosomes immediately following termination, consistent with its known function (*49, 50*).

**Fig. 2.**
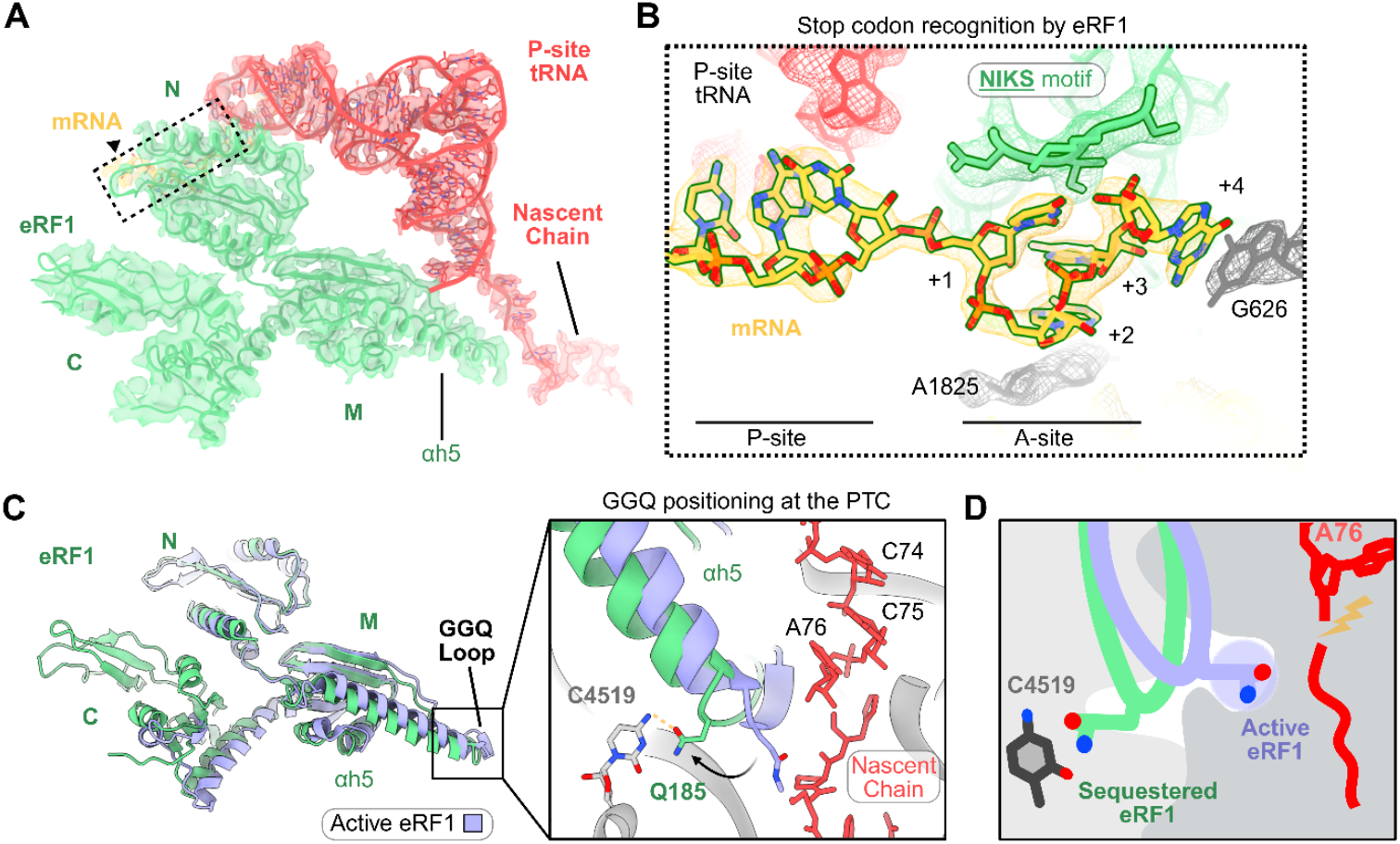
eRF1 adopts an inactive conformation on ribosomes stalled on the AMD1 C-tail. A) Overview of eRF1 domains involved in stop codon recognition (N-domain) and facilitating nascent peptide hydrolysis (M-domain). The sharpened EM map was used to visualize the N- and M-domains of eRF1 as well as the tRNA–nascent chain, while the flexible C-domain of eRF1 was visualized using a 4 Å low-pass filtered map. B) Close-up of the boxed region in (A) depicting the compaction of the stop codon in the A-site following recognition by the eRF1 N-domain. C) Comparison of eRF1 accommodation to a previous termination complex of active eRF1 (PDB 6XA1). Inset shows a closeup of the conformational changes in the GGQ loop relative to active eRF1. D) Schematic model depicting the proper positioning requirement of Gln185 for peptide hydrolysis.

Here, both eRF1 and ABCE1 natively co-purified with ribosomes paused while translating the AMD1 C-tail (Fig. 1F). The N- and M-domains of eRF1 were positioned in the A-site and resolved at 2.5-3.5 Å (fig. S2C and. S3A-E). The conserved NIKS, YxCxxxF, and GTS motifs in the eRF1 N-domain engage the A-site mRNA, and together with Gln55, coordinate stop-codon compaction and recognition as previously described (Fig. 2B, fig. S4A) (*6, 43, 44, 48*). A comparison of our structure to prior eRF1–ribosome complexes showed that the stop codon adopts the canonical architecture in the A-site following recognition (fig. S4B) (*43, 44, 48*). Notably, mutating the native stop codon to alternative stop codons (UAA/UGA) or a sense codon (GCG) did not affect ribosome stalling (fig. S4C) suggesting that stalling by the C-tail must occur prior to eRF1 binding to prevent termination.

To determine how eRF1-mediated termination and therefore nascent chain release was blocked in the ternary ABCE1:eRF1:RNC_AMD1C_ complex, we compared our structure to a previously published termination complex (*48*). Remarkably, both αh5 and the loop containing the GGQ motif undergo a conformational change which sequesters Gln185 ∼13 Å away from the peptidyl-tRNA ester bond and the PTC, in a small pocket formed by rRNA bases C4399, U4398, C4519, and C4453 (Fig. 2C, fig. S5A-B). This conformation of GGQ motif is different from recent structures of termination complexes reported from both eukaryotes and bacteria where the GGQ motif of either eRF1 or bacterial RF1 was positioned near the nascent chain to coordinate hydrolysis at the PTC (fig S5B-C) (*48, 51*). Furthermore, in our structure the P-site loop of RPL10 adopts a new conformation that contacts the tRNA, while eRF1 Phe190 stacks with A4548, which differs from the conformation observed in the active eRF1 complex (fig. S6A–B). Together, these observations show that eRF1 is sequestered and stabilized in an inactive state on the ribosome (Fig. 2D) mediated by the AMD1 C-tail arresting peptide.

### A molecular clamp orchestrates AMD1 C-tail compaction in the PET

To investigate how the C-tail is driving ribosome stalling, we analyzed its interactions with the ribosome (summarized in fig. S7A). Several turns formed by the nascent peptide, position highly conserved amino acid side chains to interact with rRNA bases that line the PET (Fig. 3A-C). Notably, Phe124, Arg121, and Lys117 (FRK clamp) capture the rRNA base A3908 by forming a series of π-stacking interactions (Fig. 3C, ‘dashed line’, Supplementary Movie 2). This clamp stabilizes the nascent chain in the tunnel before it is looped back towards the constriction site formed by uL22 and uL4. As a result, the nascent peptide adopts a unique Z-shaped configuration within the PET, accommodating an additional six amino acids in the tunnel compared with the path of a nascent peptide that does not arrest translation (fig. S8A-B). (Fig. 3A) (*52*). Furthermore, both the conformation and interactions of the stalling peptide are distinct from the structures of previously reported stalling peptides such as XBP1 (*3*) and the helical hCMV mammalian arresting peptides (*6*) (fig. S8C-D).

**Fig. 3.**
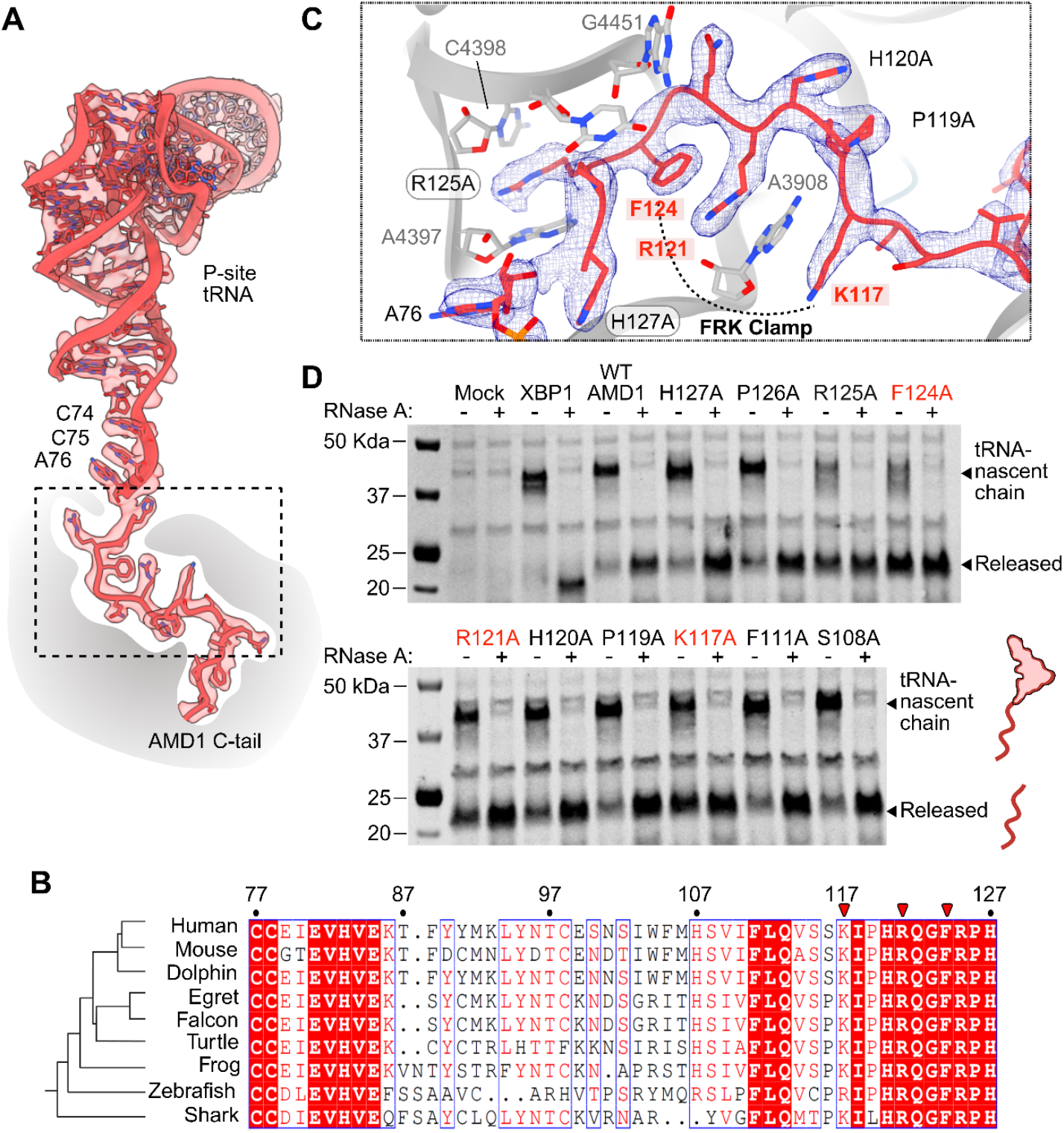
A conserved molecular clamp in the C-tail arresting peptide mediates ribosome stalling within the PET. A) Overview of the tRNA-nascent chain conformation showing the sharpened EM map as a transparent surface. B) Sequence alignment of the AMD1 C-tail among representative vertebrates. Red arrowheads indicate the clamp residues depicted in (C). C) Closeup of the early stacking interactions and molecular clamp of the AMD1 C-tail with the ribosome. The sharpened map is shown as a blue mesh. D) Ala scanning mutagenesis experiment identifying residues contributing to ribosome stalling. Clamp mutants are highlighted in red.

Sequence alignments revealed that the amino acids involved in the observed ribosome contacts were highly conserved across vertebrates (Fig. 3B). To test whether these residues were crucial for AMD1 stalling, we conducted an alanine scanning mutagenesis screen in RRL monitoring their effect on stalling relative to wild-type (WT) AMD1. Mutations of the FRK clamp significantly reduced ribosome stalling when compared to the WT AMD1 C-tail as indicated by the increased levels of released nascent chain (Fig. 3D). In contrast, mutations before and after the FRK clamp had a more subtle effect on stalling, as indicated by reduced release of the nascent chain from the tRNA and thus contribute less to ribosome stalling.

### The AMD1 C-tail inserts an arginine finger into the PTC to block translation termination

Given that the FRK clamp lies ∼15-20 Å from the peptidyl-tRNA ester bond where hydrolysis occurs, it is unclear how and whether this conformation contributes to termination inhibition. To determine specifically how translation termination was blocked and what leads to eRF1 GGQ motif sequestration away from the PTC, we closely analyzed conformational changes in the PTC triggered by the AMD1 C-tail stalling peptide. Strikingly, we found that the highly conserved Arg125 side chain of the arresting peptide inserts into the PTC where it can interfere with its catalytic activity. In particular, the Arg finger remodels the surrounding rRNA bases and sterically occludes the insertion of eRF1 Gln185, preventing it from reaching the peptidyl-tRNA ester bond where nascent chain hydrolysis occurs (Fig. 4A-C, Supplementary Movie 3). Furthermore, we show that mutation of the Arg finger alone, to a non-disruptive shorter Ala side chain, leads to nascent chain release (Fig. 4D). Together, these results reveal that ribosome stalling on the extended AMD1 translon is driven by a conserved FRK clamp, which then inserts an Arg ‘finger’ into the PTC to block eRF1 GGQ accommodation and inhibit translation termination. (Fig. 4E).

**Fig. 4.**
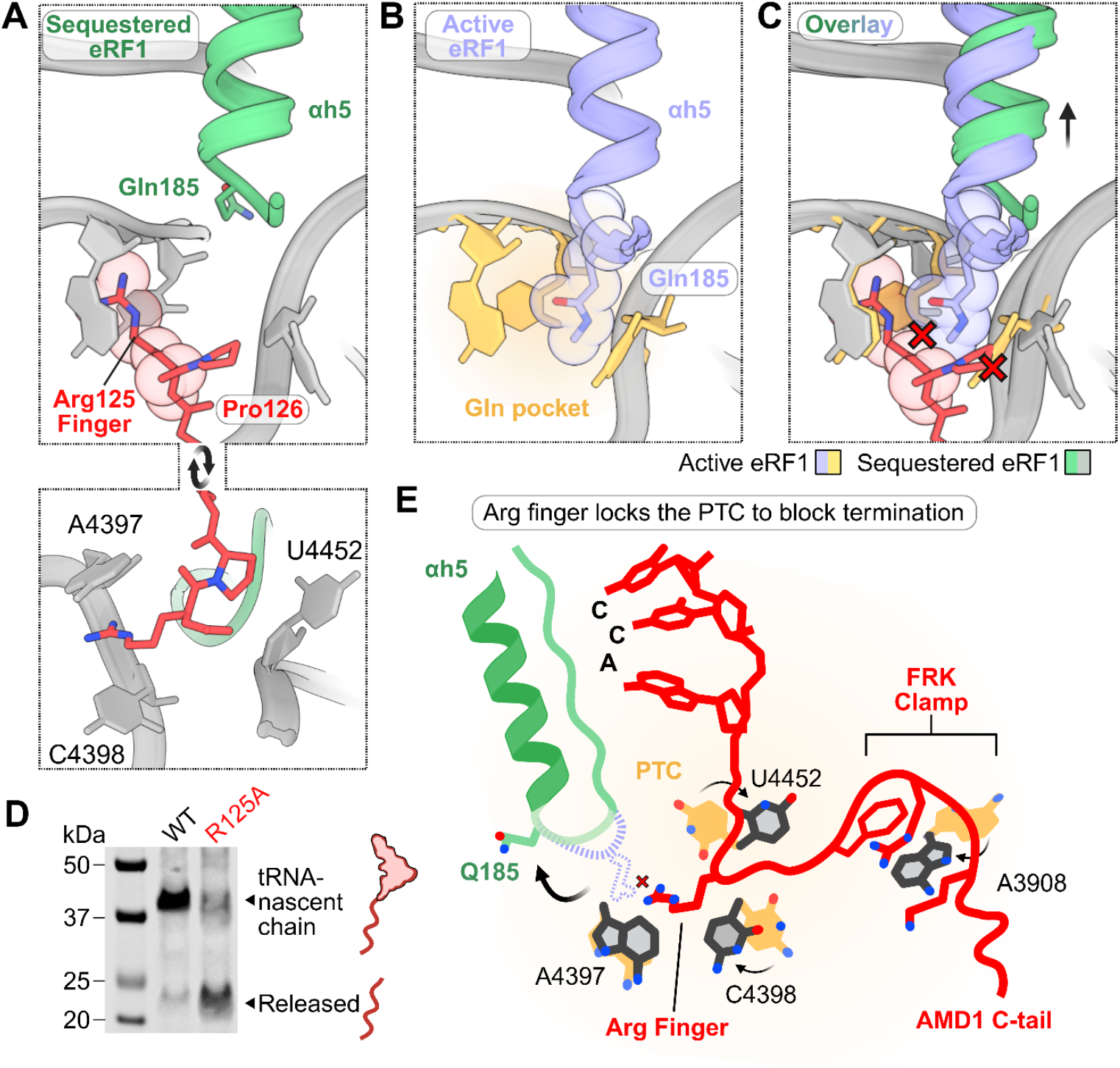
Insertion of an Arg finger into the PTC mediates termination inhibition by the AMD1 C-tail. A) Insertion of the AMD1 C-tail Arg125 side chain into the PTC. B) Proper positioning of the GGQ loop in the active eRF1 complex (PDB 6XA1). PTC residues A4397, C4398, and U4452 which coordinate positioning of Gln185 are colored yellow. C) Overlay showing the steric clash of the Arg finger with Gln185 from the active eRF1 complex. D) *In vitro* translation reaction monitoring ribosome stalling when Arg125 is mutated to Ala. Steric clashes are depicted by a red X. E) Schematic model for the mechanism of termination inhibition by the AMD1 C-tail.

### Ribosome stalling downstream of CDS in other genes

The number of available ribosome profiling samples has more than tripled since the discovery of AMD1 C-tail stalling (*39*). Aggregating these data greatly improves signal to noise ratio in ribosome profiling data analyses. Using this new aggregated dataset we manually reexamined the list of genes with high ribosome profiling density downstream of annotated CDS reported in Yordanova et al (36). For the majority of reported cases, ribosome footprint density downstream of CDS results from overlaps with translons on other transcripts. Only a few cases (e.g. *CENPA, CENPB, EEF1A2, GPX1, MACROD2, ORMDL3*) showed confident evidence of high ribosomal footprint densities likely originating from the same transcript.

To extend the search we have decided to look specifically at genes with evidence of stop codon readthrough (see Methods). After excluding cases where the peaks may have arisen due to ambiguous mapping of sequencing reads potentially originating from different loci, we identified increased peaks of density in 12 genes: *AGPAT4, AMD1, BRI3BP, CITED2, CPNE8, ETNPPL, KLC4, LDHB, MAPK10, ORMDL3, SACM1L*, and *TENT5B* (representative examples shown in fig. S8A-D). Interestingly, we found similar ribosome peaks in the orthologs of three genes i.e., *EEF1A2, MAPK10* and *SACM1L* (fig. S9A-C). Furthermore, visual examination of nucleotide conservation using a phyloP track constructed for 100-way vertebrate genomes (*53*) reveals higher conservation upstream of these stalling sites. Additionally, *MAPK10* and *SACML1* have been previously reported to have conserved readthrough regions that exhibit codon substitution patterns in multiple genomic sequence alignments, typical for protein coding genes, as measured with PhyloCSF (*54*). This suggests evolutionary selection acting on the sequences encoding nascent peptides that stall in the ribosome exit tunnel during translation, which would occur if stalling in these genes confers biological functions positively affecting Darwinian fitness. Given the sequence conservation and occurrence of ribosome stalling, it is very likely that stalling is functional not only in *AMD1*, but also in case of these three genes.

## Discussion

There is growing evidence that cis-acting elements affect translation of their own genes. *AMD1* is one such gene whose expression levels are proposed to be regulated by ribosome stalling in both its 5’ leader and 3’ trailer (*28, 36*). However, the molecular mechanism by which the C-tail induces ribosome stalling hitherto remained uncharacterized.

In this study, we revealed the molecular mechanism of ribosome stalling and termination inhibition by the AMD1 C-tail arresting peptide which halts translation at the next in-frame stop codon downstream of the *AMD1* CDS stop codon. We discovered a conserved molecular clamp in the arresting peptide that latches onto rRNA in the exit tunnel to pause translation and drive nascent chain compaction (Fig. 5). This then positions an Arg finger into the PTC altering the architecture required for GGQ loop accommodation. Thus, the AMD1 C-tail has evolved and preserved a precise strategy to lock the PTC and prevent its own release from the ribosome in a two-stage mechanism (Fig. 5). Distinct mechanisms of translational arrest have also been reported in bacteria, underscoring a broader principle in which termination inhibition is exploited across organisms to regulate translation (*7, 8, 10*).

**Fig. 5.**
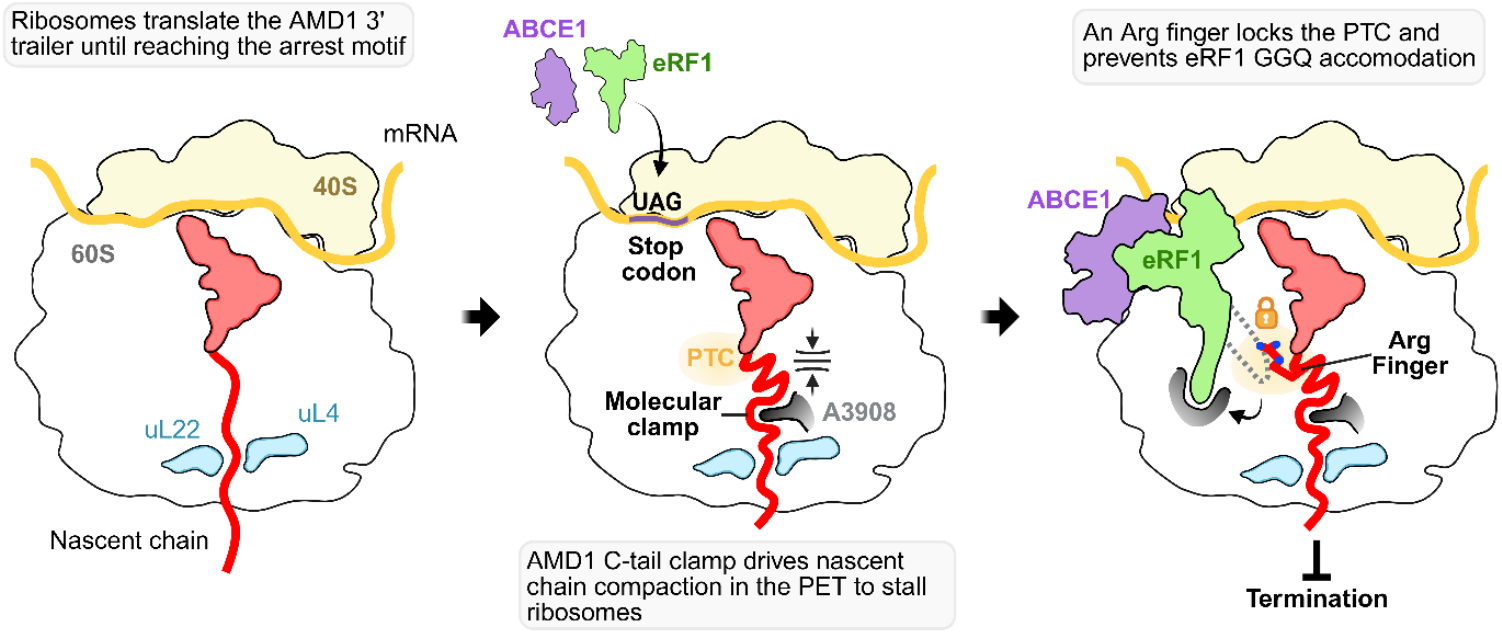
Model for ribosome stalling and termination inhibition by the AMD1 C-tail arrest peptide. Ribosomes extend into the *AMD1* 3′ trailer and pause at a downstream stop codon. The AMD1 C-tail arresting peptide forms a molecular clamp that compacts the nascent chain to stall translation. This clamp helps position an arginine side chain into the PTC, preventing eRF1 GGQ accommodation, thereby blocking translation termination.

The locked PTC state induced by the AMD1 C-tail would be also expected to perturb the positioning of any ligand in the A-site including incoming tRNAs. Notably, substituting the stop codon with a sense codon does not relieve stalling (fig. S4C), indicating that the PTC is locked and inaccessible not only to termination factors. For AMD1, the presence of a stop codon immediately following the arresting motif may provide a built-in regulatory safeguard: if ribosomes accumulate on this mRNA, termination would release the nascent chain for degradation, supporting the degron-like properties of the C-tail (*38*).

The exploration of existing ribosome profiling data revealed several human mRNA with peaks of ribosome profiling density within CDS extensions caused by potential stop codon readthrough in addition to what was reported earlier (*36*). The majority of these peaks may well result from translation of suboptimal sequences that have not been adapted to be efficiently translated due to the non-coding nature of these regions. However, we also observed that in addition to *AMD1* stalling in mRNAs of three human genes (*EEF1A2, MAPK10* and *SACM1L*) also occurs in mouse orthologs. This is accompanied by increased nucleotide conservation shortly upstream of stalling sites, suggesting the existence of purifying evolutionary selection acting on the sequence responsible for staling induction, arguing for functionality and biological significance of these stalls.

The mechanism of PTC locking that couples ribosome stalling with termination evasion by the conserved AMD1 C-tail arresting peptide, carries broader implications for how mRNA translation can be modulated through ribosome–nascent peptide interactions. What cellular signals trigger ribosome pausing in the 3′ UTR of *AMD1* and other genes with analogous stalling motifs and how these stalled ribosomes are rescued remain important questions for future investigation.

## Acknowledgments

We thank Pramod Bhatt on early discussions of AMD1 construct design and Aashritha Penumudi for help with the ribosome stalling and purification experiments. Cryo-EM data was collected at the Molecular Electron Microscopy Core (RRID:SCR_019031) established at the University of Virginia (UVA) School of Medicine with support from the National Institutes of Health (NIH) grants G20-RR31199, S10-RR025067, and U24-GM116790. We thank Michael Purdy for his support with cryo-EM data collection.

## Author contributions

E.M.: conceptualization, data curation, formal analysis, investigation, methodology, visualization, writing- original draft, writing-review and editing; F.S.B.: formal analysis, methodology, investigation, funding acquisition; M.I.S.: formal analysis, methodology, investigation; K.A.M.: methodology; M.M.Y.: conceptualization, supervision, writing-review and editing; P.V.B.: conceptualization, funding acquisition, project supervision, writing- review and editing; A.J.: conceptualization, funding acquisition, project supervision, writing- review and editing. All authors contributed to data analysis and the final version of the manuscript.

## Data and materials availability

Cryo- EM maps and model coordinates are deposited in the Electron Microscopy Data Bank (EMDB) and the Protein Data Bank (PDB) under accession codes: EMD-XXXX and PDB-YYY. All other data are available in the main text of the paper or in the supplementary material.

