## Supplemental Figures for "Mechanism of Ribosome Stalling by the AMD1 C-terminal Tail Arrest Peptide"

**The PDF file includes:**

Materials and Methods
Figs. S1 to S9
Table S1
References

### Materials and Methods

Cloning and Plasmid Preparation

Constructs used in this study were designed as previously described (1). Sequences encoding for *AMD1 (AAs 78-127)* and *XBP1 (AAs 237-261)* were obtained as two gBlocks (Integrated DNA Technologies, Belgium). Both sequences included, from 5′ to 3′, a T7 promoter, 3×FLAG epitope, perfect Kozak sequence, flexible linker, and the indicated stalling sequence.

The *AMD1* gBlock was cloned into a PCR-linearized pUC19 vector using Gibson assembly (2) with a home-made mastermix. Site-specific mutagenesis was performed using In Vivo Assembly (IVA) cloning (3), in which all candidate codons were substituted with GCG (alanine). All PCR amplifications were carried out with Q5 High-Fidelity DNA Polymerase (New England Biolabs, Cat. No. M0491L). For IVA cloning, DpnI (New England Biolabs, Cat. No. R0176L) digestion was performed to remove parental DNA templates.

All constructs were transformed into *E. coli* DH5α competent cells and purified using the GeneJET Plasmid Miniprep Kit (Thermo Fisher Scientific, Cat. No. K0503). Correct sequences were confirmed by Sanger sequencing (Eurofins Genomics, Germany).

*In vitro* Transcription

PRC fragments encoding a T7 promoter and a 3X-FLAG tag linked to the AMD1 C-tail (AAs 78-127) were *in vitro* transcribed for 4 hours at 38°C with 1.1 mg/mL T7 polymerase in reaction buffer (40 mM Tris-HCl pH 7.6, 5 mM ribonucleoside triphosphates, 6 mM $MgCl_2$, 2 mM spermidine, 1 mM DTT, 0.04 U/uL RNase Inhibitor). Precipitate was pelleted at 14,000 rpm for 5 minutes at 4°C. The resulting supernatant was placed in a sterile tube and one volume of 6M LiCl was added. The mixture was incubated for 1 hr at 4°C and then centrifuged at 14,000 rpm for 20 minutes at 4°C. The supernatant was discarded, and the resulting pellet was washed with 200 µL of ice-cold 70% ethanol and centrifuged at 14,000 rpm for 5 minutes at 4°C. The supernatant was discarded, and the pellet was resuspended in 100 µL of sterile $ddH_2O$. Resuspended RNA was placed on ice for 5 minutes and 40 µL of 2.8 M NaOAc and 300 µL of ice-cold 95% ethanol were added. The mixture was incubated on ice for an additional 5 minutes and then centrifuged at 14,000 rpm for 30 minutes at 4°C. The resulting pellet was washed with 200 µL of ice-cold 70% ethanol and centrifuged at 14,000 rpm for 5 minutes at 4°C. After discarding the supernatant, the purified RNA pellet was resuspended in 100 µL of sterile $ddH_2O$. The RNA concentration was determined using a nanodrop (ThermoFisher).

*In vitro* Translation Reactions for Monitoring Ribosome Stalling

The mRNA corresponding to the AMD1 variants and XBP1 control were *in vitro* translated in rabbit reticulocyte lysates (RRL) diluted to 66.7% (v/v) in buffer (0.5 µg/µL mRNA, 81 mM KCl, 2 mM $Mg(OAc)_2$, 24 µM amino acid mix, 0.2 µM spermidine, 0.04 U/µL RNase Inhibitor (ThermoFisher), 0.5X protease inhibitor (Promega)). The reactions were conducted at 32 °C for 25 minutes and then moved to ice. Samples were diluted in SDS-PAGE loading dye (final concentration: 50 mM Tris-HCl pH 6.8, 2% SDS (w/v), 143 mM β-mercaptoethanol, 6% glycerol (v/v), 0.004% bromophenol blue (w/v)) and loaded on a 4-12% Bis-Tris gel (Genescript, cat no. M00654). The remaining sample was flash frozen in liquid nitrogen.

Proteins were resolved and then transferred onto a 0.2 μm nitrocellulose membrane (LI-COR, cat no. 926-31092). Membranes were blocked in 5% milk in 1X PBST (0.1% Tween-20) for 1 hr at RT, with shaking. The membranes were incubated with primary antibodies diluted in 2% BSA in 1X PBST for 1 hr at RT or overnight at 4°C, with shaking. Primary antibodies: α-FLAG (Sigma, cat no. F1804, 1:3000), α-RPL10a (Invitrogen, cat no. MA5-44710, 1:2000). Membranes were incubated for 1 hr at RT with secondary antibodies diluted in 5% milk in 1X PBST. Secondary antibodies used: α-mouse (Invitrogen, cat no. A21058, 1:10,000) and α-rabbit (Invitrogen, cat no. A32735, 1:10,000). Membranes were imaged with the LI-COR odyssey imager.

Alanine scanning mutagenesis experiments

Mutant variants of the AMD1 C-tail were PCR-amplified and mRNA was *in vitro* transcribed and purified using the RNA Clean & Concentrator Kit (Zymo, cat no. R1016). The resulting mRNA was translated as described above or using a homemade RRL as previously described (1). Briefly, 20 μL rabbit reticulocyte lysate (RRL) reactions were prepared. Each translation reaction was split into two equal aliquots: one was treated with RNase A (Promega, Cat. No. A7973) for 20min at 30°C, while the untreated was left in ice. From each aliquot, 1 or 2 μL were diluted in SDS-PAGE loading dye (final concentration: 125 mM Tris-HCl pH 6.8, 4% SDS (w/v), β-mercaptoethanol 10% (v/v), 20% glycerol (v/v), 0.004% bromophenol blue (w/v)), loaded into a Bolt™ Bis-Tris Plus 4–12 % gradient mini gels, WedgeWell™ format (Invitrogen, Cat. No. NW04127BOX) and run for 23 minutes. Proteins were transferred to nitrocellulose membranes for 7 minutes using a Trans-Blot Turbo™ Transfer System (Bio-Rad). Membranes were blocked in 5 % BSA in PBST (0.1 % Tween-20) for 1 h at room temperature, followed by incubation with primary antibody anti-FLAG (Sigma, Cat. No. F1804-200UG; 1:1000 in 1 % BSA in PBST) overnight at 4°C with gentle shaking. After washing, membranes were incubated with secondary antibody IRDye® 800CW Donkey anti-Mouse IgG (H+L) (Cat. No. 926-32212; 1:20,000 in 1 % BSA in PBST) for 30mins at room temperature. Signals were visualized using a LI-COR Odyssey imaging system.

$\underline{\text{RNC}_{\text{AMD1C}}}$ Purification

RNCs were purified as previously described (*4*). mRNA was *in vitro* translated using RRL (Promega, cat no. L4540) diluted to 66.7% (v/v) in buffer (0.5 μg/μL mRNA, 81 mM KCl, 2 mM $Mg(OAc)_2$, 24 μM amino acid mix, 0.2 μM spermidine, 0.04 U/μL RNase Inhibitor (ThermoFisher), 0.5X protease inhibitor (Promega)) at 32 °C for 25 minutes and then moved to 4°C. The reaction was incubated with a 200 μL slurry of Anti-DYKDDDK beads (Sigma, cat no. A2220) for 2 hours at 4°C with rotation in a gravity column. Following incubation, the beads were pooled to the bottom of the column using 650 μL of high-salt wash buffer (50 mM HEPES-KOH pH 7.7, 750 mM KOAc, 10 mM $Mg(OAc)_2$, 1 mM DTT, 0.1% Triton-X). The column was washed with 10 bead volumes of high-salt wash buffer twice followed by two washes with 10 bead volumes of low-salt wash buffer (50 mM HEPES-KOH pH 7.7, 100 mM KCl, 10 mM $MgCl_2$). Elutions were carried out at room temperature with 110 μL of 0.25 mg/mL FLAG peptide in ribosome buffer (50 mM HEPES-KOH pH 7.7, 100 mM KOAc, 15 mM $Mg(OAc)_2$) for 15 minutes (5X). Eluted fractions were placed on ice, combined, and then centrifuged at 100,000 rpm for 1 hr at 4 °C using the TLA120.1 rotor (Beckman). The supernatant was discarded, and the pellet was resuspended in a ribosome buffer using a micropipette. The concentration of the purified ribosomes was determined with the nanodrop.

Cryo-EM sample preparation and data acquisition

Immediately following purification, RNCs were diluted to ~250 ng/µL in ribosome buffer and placed on ice. Quantifoil R2/1 300 mesh copper grids (Quantifoil, cat no. Q350CR1) were coated with a thin layer of 3.3 nm carbon. The grids were glow discharged at 15 mA for 15 seconds (EMS). A 5 µL sample of the purified RNCs was incubated on the grid for 1 min at 4°C and 100% relative humidity. Following the incubation period, grids were blotted and plunged into liquid ethane pre-cooled to liquid nitrogen temperatures using a Vitrobot Mark IV (ThermoFisher). Data was collected at the University of Virginia Molecular Electron Microscopy core using the ThermoFisher Krios electron microscope operated at 300 kV and equipped with a K3 direct electron detector at a nominal magnification of 105,000x (pixel size of 0.83 Å), with an energy filter slit width of 10 eV. Data was collected at a defocus range of -1.8 to -0.8 µm with a step size of 0.2 µm. A dose of 50 e/$Å^2$ was applied across 40 frames per movie and a total of 11,162 movies were collected.

Single particle cryo-EM data processing

Cryo-EM data was processed using CryoSPARC v4.6.0 (*5*). Movies were patch motion corrected and subjected to patch CTF estimation. Particles with minimum and maximum diameters of 250 Å and 400 Å, respectively, were selected using blob picker. Particles were extracted at a box size of 464 x 464 pixels and binned to 128 pixels for initial 2D and 3D classifications. The 173,341 particles from the 2D classification showing ribosome classes were used for ab-initio reconstruction to generate initial models for subsequent heterogeneous refinement to remove rotated and collided ribosomes. The particles were then re-extracted at full box size without binning and refined to generate a consensus ribosome structure at 2.76 Å. Iterative 3D variability analyses (*6*) were conducted to sort for a homogenous stack of particles containing high resolution features for the additional factors observed in the consensus structure. Focused masks were used encompassing the A- and P-site of the ribosome. Following three rounds of 3D variability analyses, a final set of 15,310 particles containing the bound factors were refined to an average resolution of 2.86 Å determined by the gold standard Fourier Shell Correlation (FSC) with an FSC cutoff of 0.143. The local resolution was calculated in CryoSPARC at an FSC threshold of 0.143.

Model Building

A previously published models of the 80S ribosome (PDB 6R5Q), eRF1 (PDB 5LZU), and ABCE1 (PDB 3JAH) were fitted into the final cryo-EM map using ChimeraX. Individual domains and helices of eRF1 and ABCE1 were further adjusted as rigid bodies in Coot v0.9.6 (7). Models were manually adjusted based on observed side chain densities in the EM map. The AMD1 C-tail arresting peptide atomic model was built *de novo* based on the observed side chain densities using available sequence information. Due to the lack of a reliable sequence for the mammalian tRNA-His (AUG), the tRNA-His sequence from *Danio rerio* (tRNA-His-ATG-3-1 from GtRNAdb) was used to model the P-site tRNA. This sequence largely matched the EM density with one A/U and G/C base swap. The final model was iteratively refined in PHENIX v1.20.1-4487 using 3 macrocycles of real space refinements applying Ramachandran, sidechain rotamer, protein secondary structure and nucleotide restraints (*8*). Molprobity was used to validate the model and resolve clashes. All cryo-EM figures and models were made in ChimeraX.

Ribosome profiling data analysis

For ribosome profiling data analysis, we used 3835 human libraries from RiboSeq Data Portal (9) mapped to the human genome in RiboCrypt. For the analysis of *AMD1* ortholog: species supported by RiboCrypt with at least 3 ribosome profiling libraries were used to generate *AMD1* transcript-level ribosome occupancy plots (Fig. S1). For the analysis of pausing at the readthrough extensions we have included all cases annotated as readthrough according to GENCODE v.47 catalogue except for *AGO1*, *VEGFA* and *ACP2* that are lacking ribosome profiling support for stop codon readthrough in RiboCrypt upon manual examination. In addition, we identified candidate stop codon readthrough genes based on the following parameters using ORFik R library (10) (extension length >6 codons, ORFScore: > 5, In-frame RPF reads: > 300, Codons covered: > 3, Codons with in-frame reads: > 10%, In-frame vs. out-of-frame read ratio: > 50%). Subsequently any cases where readthrough extensions overlap annotated CDS from other transcripts have been excluded. The resulting 78 readthrough candidates have been examined for the presence of outlier peaks of ribosome profiling density at individual positions. A peak of read density was considered to be an outlier if its density Z-score >5. Subsequently the profiles containing such peaks have been examined manually. For the 12 cases with confirmed outlier peaks we have analyzed nucleotide conservation using phyloP. We also have examined ribosome profiles of mouse orthologs for these candidates.

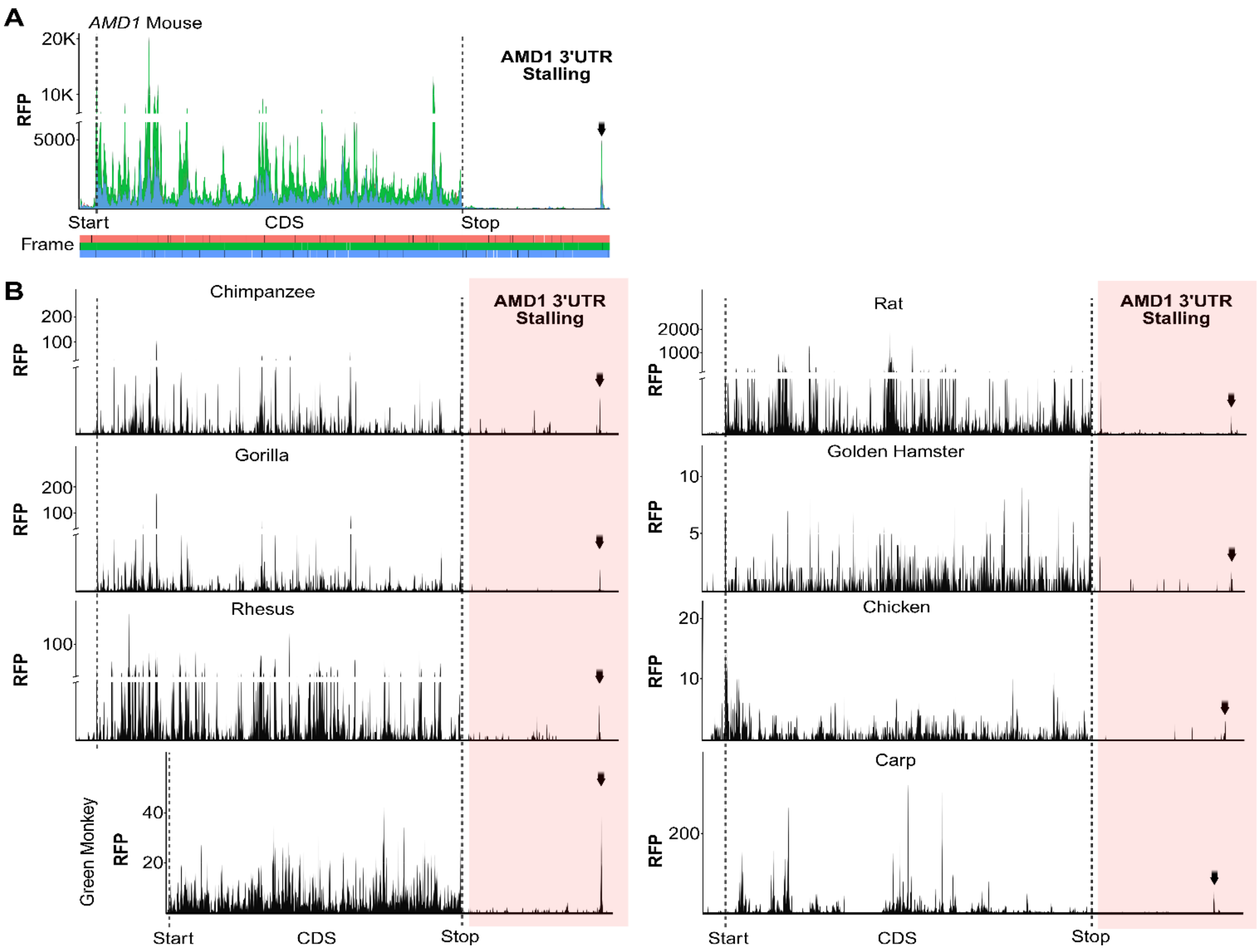

**Fig. S1**

***AMD1* ribosome profiles of vertebrate homologs.** A) Aggregated ribosome sequencing reads for the mouse *AMD1* transcript (ENSMUST00000099945) showing the CDS and a part downstream of it that contains the sight of ribosome stalling. B) Ribosome profiles of *AMD1* mRNAs for the indicated vertebrates highlighting peaks present at the end of *AMD1* extended translon (red). The green monkey (*Chlorocebus sabaeus*) ENSEMBL transcript ENSCSAT00000013631.1 is truncated according to our examination of the genomic region and is missing two 5' exons.

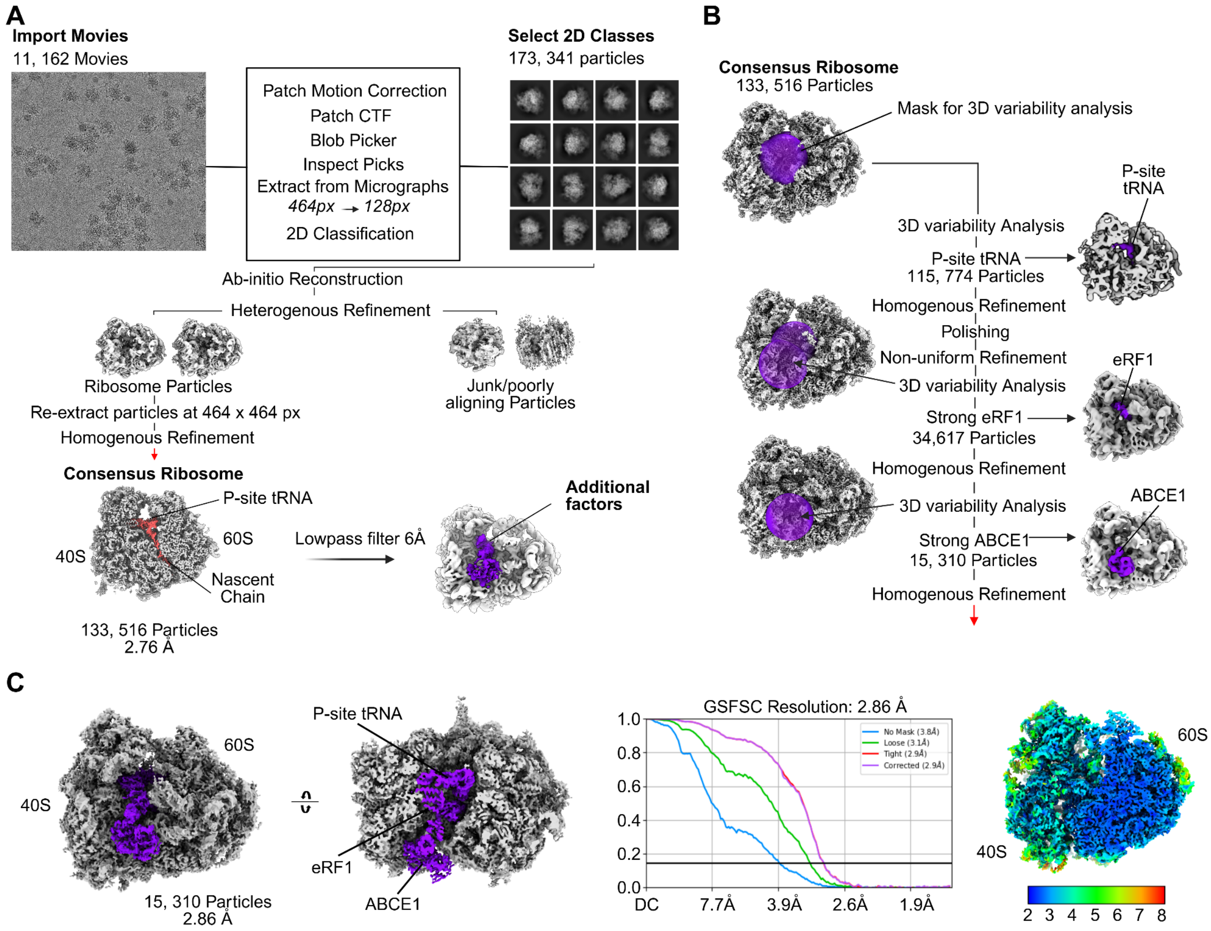

**Fig. S2**

**Single particle cryo-EM processing workflow used to determine the ABCE1:eRF1:RNC$_{\mathbf{AMD1C}}$ structure.** A) Movies were patch motion corrected and CTF estimated in CryoSPARC. Blob picker was used to select for particles from the resulting corrected micrographs. Particles were extracted at a box size of 464 x 464 pixels and binned to 128 pixels. Following 2D classification, 173, 341 particles were selected and used to generate ab-initio models. The models were used in subsequent heterogenous refinements to sort for ribosome particles and discard junk or poorly aligning particles. Consensus ribosomes were homogeneously refined and displayed strong P-site and nascent chain density (red) along with additional density for factors interacting at the A-site of the ribosome (purple). B) The consensus ribosome particles were iteratively classified by 3D variability analysis using the indicated focused masks (purple spherical mask) to reach a homogenous subset of ribosome particles containing P-site tRNA, eRF1 and ABCE1 (purple). C) The final eRF1:ABCE1:RNC$_{AMD1C}$ particles were homogeneously refined to an average resolution of 2.86 Å, determined by gold standard Fourier Shell Correlation (GSFSC) at an FSC cutoff of 0.143. The local resolution was estimated in cryoSPARC at an FSC threshold of 0.143.

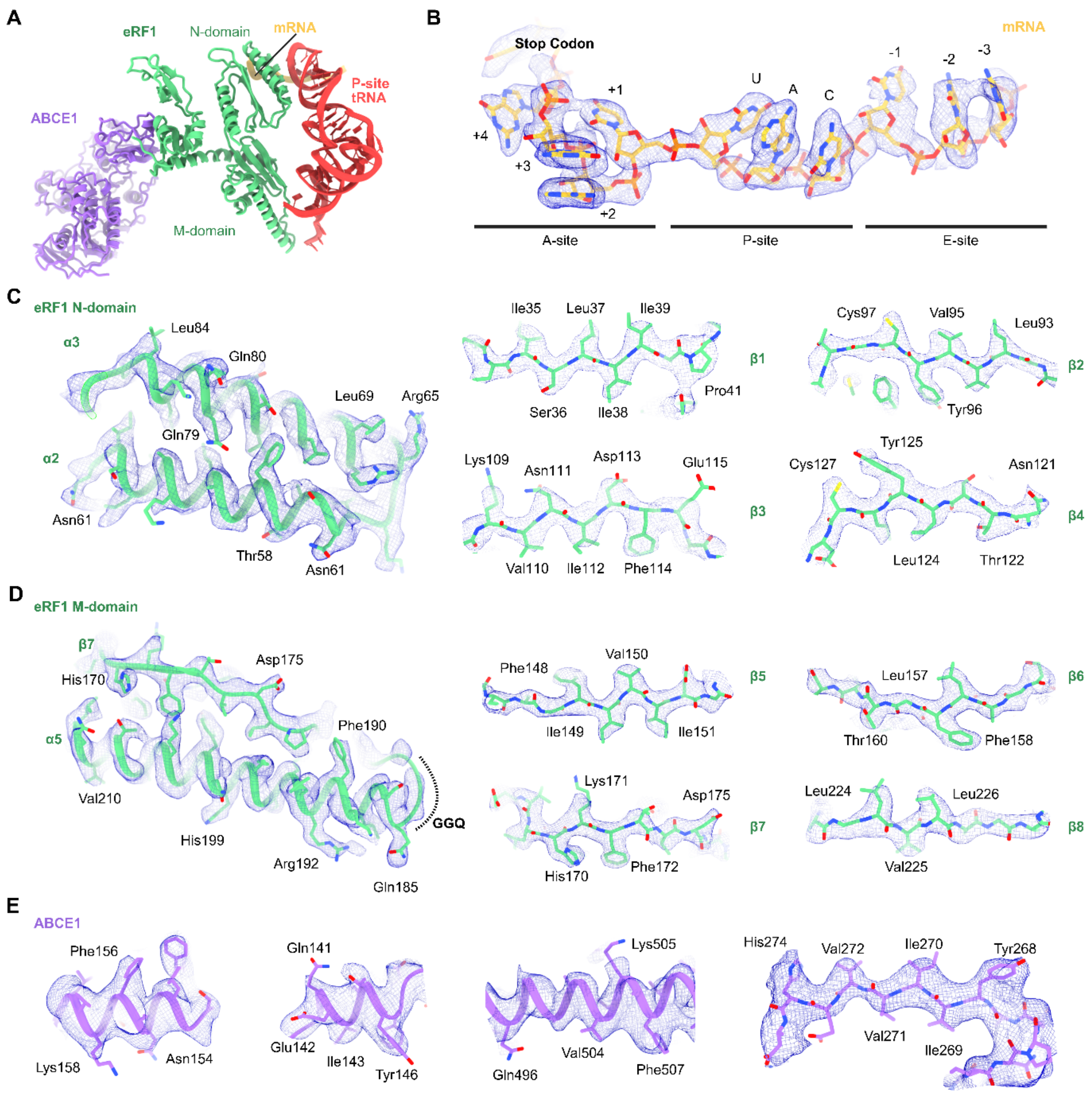

**Fig. S3**

**Structure of ABCE1 and eRF1 in complex with a P-site tRNA and mRNA on the stalled ribosome.** A) Overview of the ABCE1, eRF1, mRNA, and P-site tRNA models shown without the rest of the ribosome for clarity. B-E) Fits for the indicated model regions of the mRNA, eRF1 N- and M-domains, and ABCE1. Sharpened EM maps are used to visualize the mRNA and eRF1. Unsharpened maps were used for visualizing ABCE1.

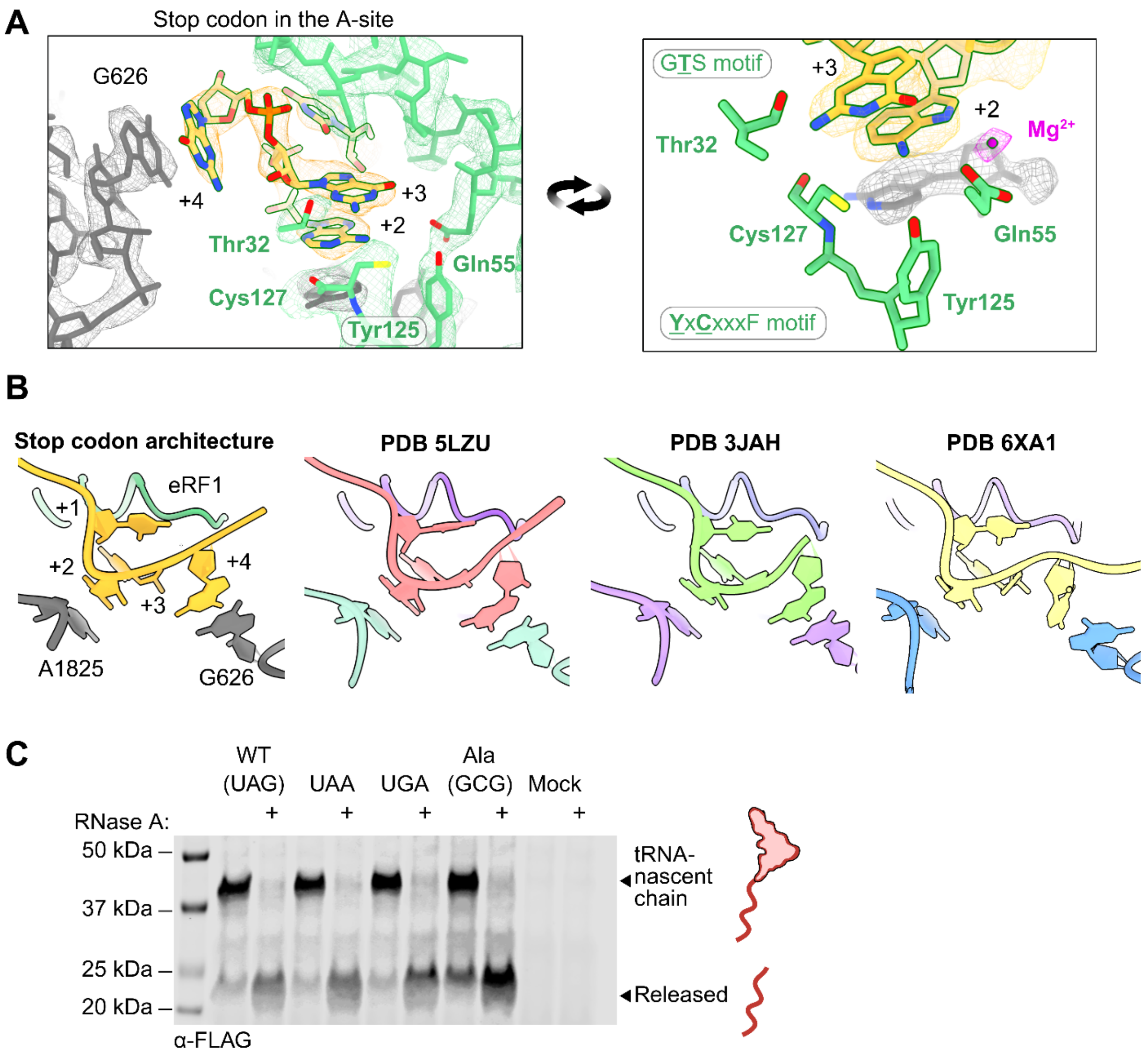

**Fig. S4**

**Stop codon recognition in the A-site by eRF1 does not affect ribosome stalling.** A) Closeups of the stop codon interactions with the N-domain motifs of eRF1. Sharpened maps were used for visualization. B) Comparison of stop codon compaction following recognition by eRF1 with other termination complexes (PDB 5LZU, 3JAH, 6XA1). C) Mutations of the native stop codon in the AMD1 C-tail.

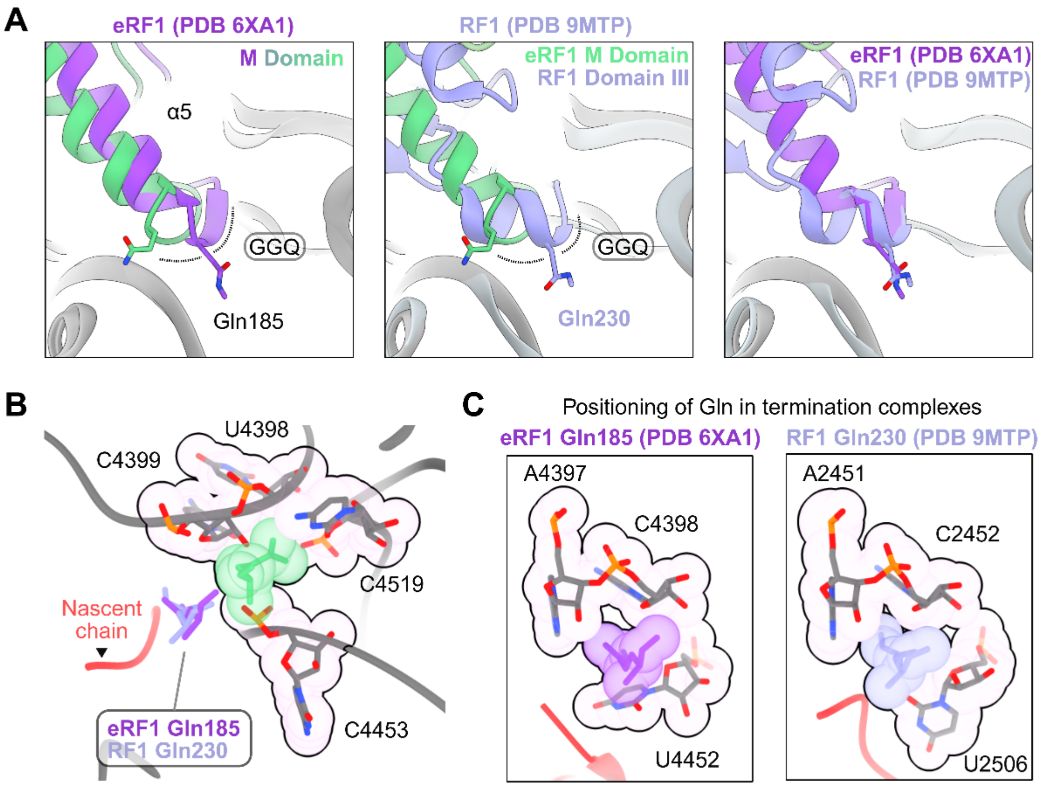

**Fig. S5**

**Comparison of GGQ motif in AMD C-tail stalled ribosome versus other termination complexes.** A) Comparison of GGQ positioning in the current structure (green) and other termination complexes (purple) of eRF1 and RF1. B) Sequestration of the eRF1 Gln185 (green) on ribosomes stalled by the AMD1 C-tail in a pocket away from the PTC compared to other termination complexes (purple). C) Closeups of the proper positioning of glutamines that make up the GGQ motif at the PTC in other termination complexes. Models used for comparison: PDB 6XA1 (human eRF1), PDB 9MTP (bacterial RF1).

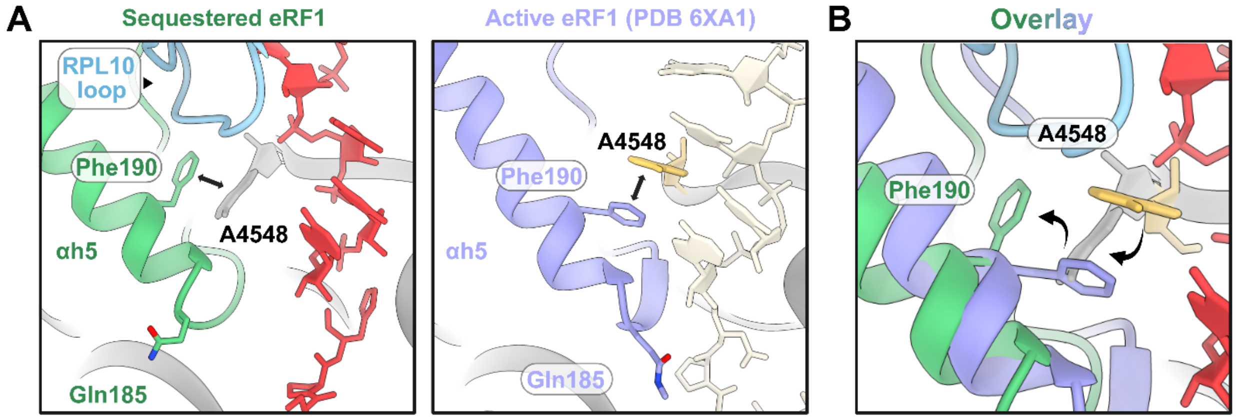

**Fig. S6**

**A conformational change in eRF1 αh5 and RPL10 interactions observed in the AMD C-tail stalled ribosome.** A) The flexible loop of RPL10 was resolved interacting with the P-site tRNA. eRF1 Phe190 π-stacks with A4548 in a distinct conformation when sequestered. B) Closeup of the conformational changes in Phe190 and A4548 relative to the previous termination complex (PDB 6XA1).

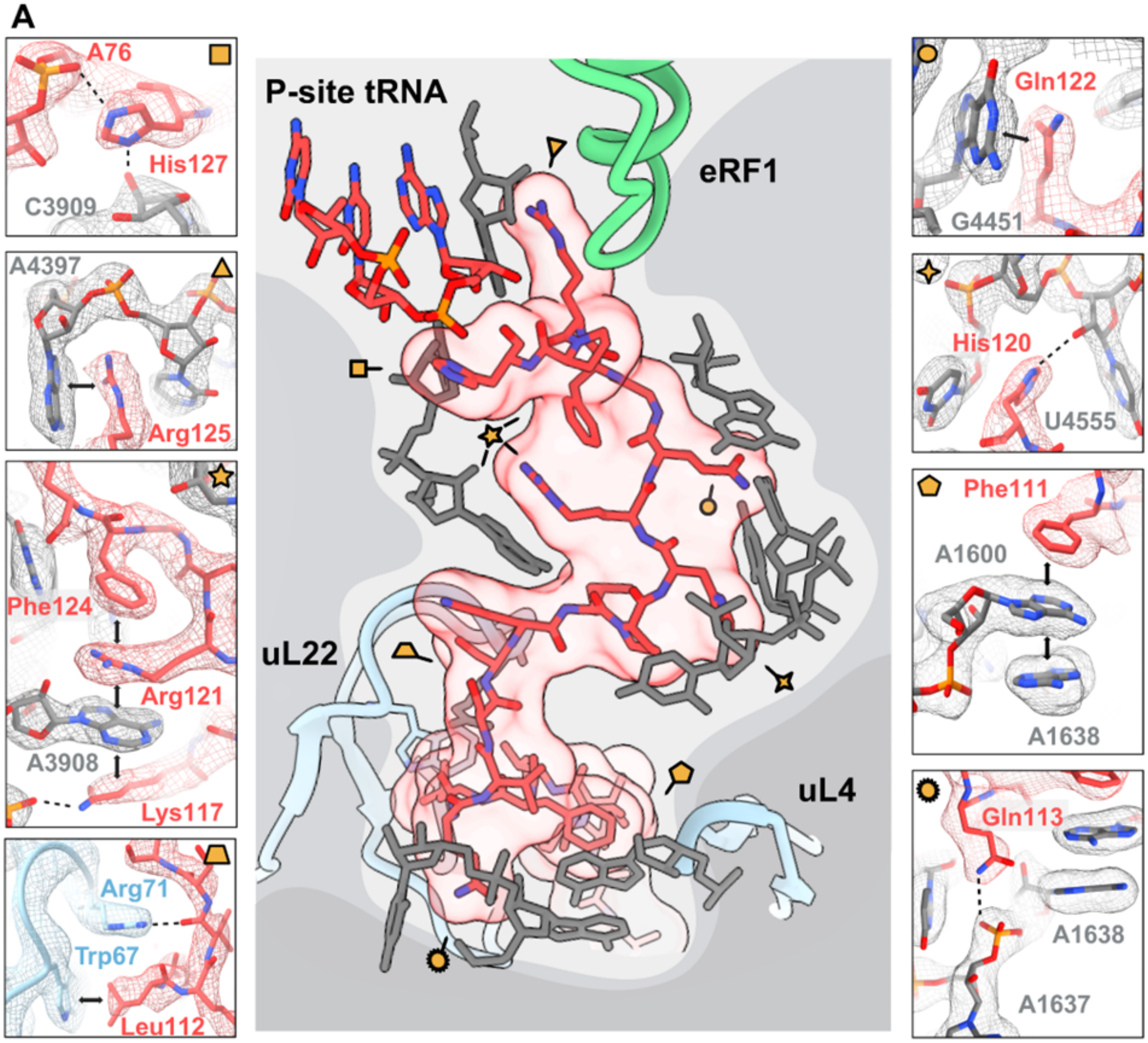

**Fig. S7**

**The AMD1 C-tail stalls the ribosome via an extensive interaction network in the exit tunnel.**

A) Central panel depicts a model of the nascent chain shown as a transparent surface flanked by interacting bases and ribosomal proteins from the PET. From the top left: hydrogen bond between His127 and the backbone of A76 and C3909; π-stacking interaction between Arg125 and A4397; π-stacking network formed between Phe124, Arg121, A3908, and Lys117; hydrogen bond formed between Arg71 (uL22) with the carbonyl carbon of Val19 and hydrophobic interaction between Trp67 (uL22) and Leu112. From the top right: the AMD1 C-tail forms an additional π-stacking interaction between Gln122 and G4451; polar interaction between His120 and U4555; π-stacking interaction between Phe111, A1600, and A1638; polar interaction between Gln113 and A1638 backbone. Sharpened maps are depicted as mesh for visualization.

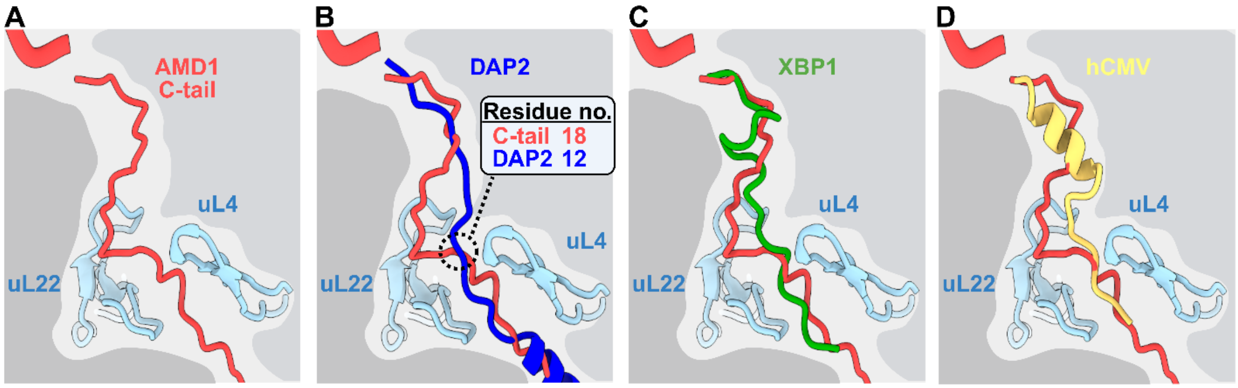

**Fig. S8**

**Comparison of nascent chains in the exit tunnel of mammalian arresting peptides and the non-arresting peptide DAP2 in the PET.** A) The AMD1 C-tail adopts a unique Z-shaped configuration in the exit tunnel leading to the constriction site formed by uL22 and uL4. B) DAP2 does not display any notable compaction or secondary structural features when compared to the AMD1 C-tail. C) XBP1 adopts a short, compacted S-shaped configuration when compared to the AMD1 C-tail. D) The hCMV arresting peptide forms a short α-helix to stall ribosomes while the AMD1 C-tail does not form any secondary structures prior to the constriction site. Models used for comparison: DAP2 (PDB 7OBR), XBP1 (PDB 6R5Q), hCMV (PDB 5A8L).

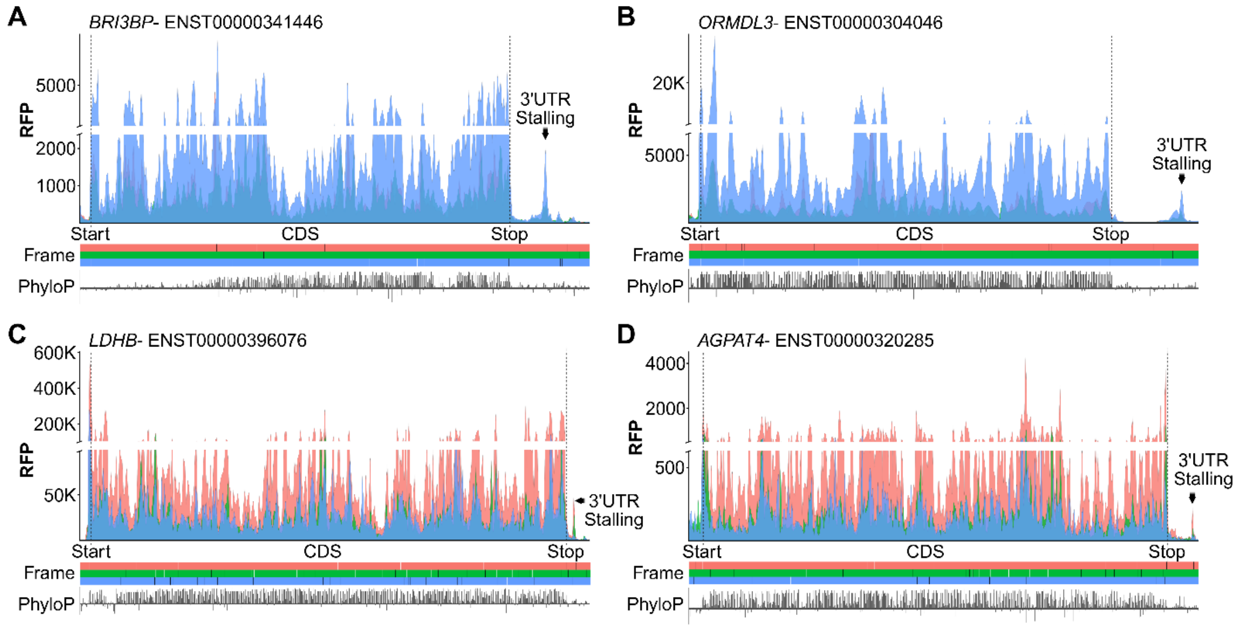

**Fig. S8**

**Representative ribosome profiles for the indicated human genes with stalling present in the 3'UTR.** Aggregated human sequencing reads for *BRI3BP* (A), *ORMDL3* (B), *LDHB* (C), and *AGPAT4* (D). Plots were generated with ribocrypt.org and show the CDS region with potential stop codon readthrough regions containing stalling sites as indicated.

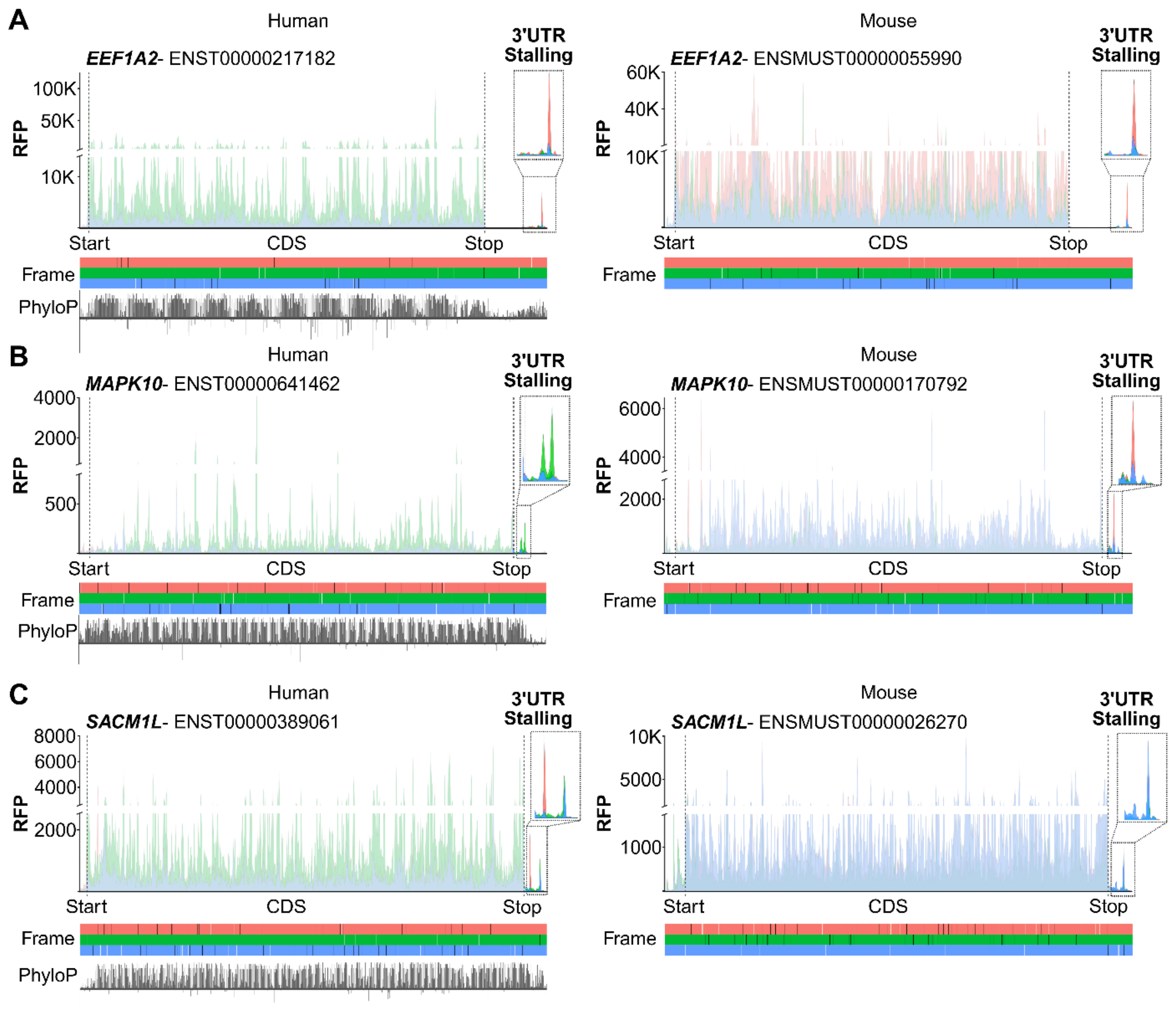

**Fig. S9**

**Ribosome stalling in 3' trailers of select candidates present in mice ribosome sequencing data.** A) Ribosome profile of human (left) and mouse (right) *EEF1A2*. B) Ribosome profile of human (left) and mouse (right) *MAPK10*. C) Ribosome profile of human (left) and mouse (right) *SACM1L*. All plots are aggregated data of ribosome sequencing reads from multiple studies generated with ribocrypt.org showing the CDS region (transparent) with indicated stalling sites at the end of extended translons.

| | eRF1:ABCE1:AMD1$_{C\text{-}tail}$ |
|---|---|
| EMDB code | EMD-XXXX |
| PDB code | YYYY |
| **Data collection and processing** | |
| Nominal magnification | 105,000x |
| Voltage (kV) | 300 |
| Electron exposure ($e^{-}$/Å$^2$) | 50 |
| Defocus range (μm) | -1.8/-0.8 |
| Pixel size (Å) | 0.83 |
| Initial particle images (no.) | 133, 516 |
| Final particle images (no.) | 15, 310 |
| Map resolution at FSC=0.143 (Å) | 2.86 |
| **Structure refinement in PHENIX 1.20.1** | |
| Model resolution at FSC=0.5 (Å) | 3.1 |
| $CC_{mask}$ | 0.80 |
| Map sharpening B factor (Å$^2$) | 29 |
| Model composition | |
| Non-hydrogen atoms | 218669 |
| Protein residues | 12612 |
| RNA residues | 5472 |
| B factors min/max/mean (Å$^2$) | |
| Protein | 0.00/109.15/52.60 |
| RNA | 0.00/165.06/63.53 |
| Ligand | 0.00/125.13/27.89 |
| RMSD | |
| Bond lengths (Å) | 0.002 |
| Bond angles (°) | 0.451 |
| **Validation** | |
| MolProbity score | 1.65 |
| Clashscore | 8.53 |
| Poor rotamers (%) | 1.09 |
| Ramachandran plot | |
| Favored (%) | 0.08 |
| Allowed (%) | 2.85 |
| Outliers (%) | 97.07 |

**Table S1.**
**Cryo-EM data collection, model refinement and validation statistics.**
